# Single-cell Visualization and Quantification of Trace Metals in Chlamydomonas Lysosome-Related Organelles

**DOI:** 10.1101/2021.02.12.431000

**Authors:** Stefan Schmollinger, Si Chen, Daniela Strenkert, Colleen Hui, Martina Ralle, Sabeeha S. Merchant

## Abstract

The acidocalcisome is an acidic organelle in the cytosol of eukaryotes, defined by its low pH and high calcium and polyphosphate content. It is visualized as an electron-dense object by transmission electron microscopy (TEM) or described with mass-spectrometry (MS)-based imaging techniques or multimodal X-ray fluorescence microscopy (XFM) based on its unique elemental composition. Compared to MS-based imaging techniques, XFM offers the advantage of absolute quantification of trace metal content, since sectioning of the cell is not required and metabolic states can be preserved rapidly by either vitrification or chemical fixation. We employed XFM in *Chlamydomonas reinhardtii*, to determine single-cell and organelle trace metal quotas within algal cells in situations of trace metal over-accumulation (Fe, Cu). We found up to 70% of the cellular Cu and 80% of Fe sequestered in acidocalcisomes in these conditions, and identified two distinct populations of acidocalcisomes, defined by their unique trace elemental makeup. We utilized the *vtc1* mutant, defective in polyphosphate synthesis and failing to accumulate Ca to show that Fe sequestration is not dependent on either. Finally, quantitation of the Fe and Cu contents of individual cells and compartments via XFM, over a range of cellular metal quotas created by nutritional and genetic perturbations, indicated excellent correlation with bulk data from corresponding cell cultures, establishing a framework to distinguish the nutritional status of single cells.

**Significance statement:** Transition metals are of crucial importance for primary productivity; their scarcity limits crop yield in agriculture and carbon sequestration at global scale. Copper (Cu), iron (Fe) and manganese (Mn) are among the most important trace elements that enable the redox chemistry in oxygenic photosynthesis. The single-celled, eukaryotic green alga *Chlamydomonas reinhardtii* is a choice experimental system for studying trace metal homeostasis in the context of phototrophy, offering all the advantages of a classical microbial system with a well-characterized photosystem and trace metal metabolism machinery of relevance to plants. This project identifies and differentiates different trace metal storage sites in Chlamydomonas and uncovers the dynamics of trace metal storage and mobilization in situations of fluctuating resources.

## Introduction

Trace metals like iron (Fe), copper (Cu), manganese (Mn) and zinc (Zn) play a crucial role as co-factors, providing a range of chemical capabilities to enzymes central to life. Metals also provide structural stability to proteins (1, 2) and enable the catalysis of essential metabolic reactions by providing functional groups that are not readily available via the side chains of amino acids. Consequently, trace metals are required in ∼ 40 % of all enzymes as part of their catalytic centers (3, 4). Individual enzymes are often optimized to utilize a specific metal cofactor, with respect to the chemical utility of that particular trace metal for the catalyzed reaction, the metal’s specific requirements for the binding site in the protein (dimensions, charge, coordination preferences), and the availability of the trace metal within the organism’s reach. There is, however, the possibility of mis-metalation, largely attributed to protein structural flexibility and somewhat similar ionic radii and coordination preferences of first-row trace metals (5, 6). Enzyme mis-metalation can harm the cell directly by loss-of-function (7, 8), by accumulation of unintended products, or the production of toxic side products, for example reactive oxygen species (9). Cells have therefore developed elaborate strategies to facilitate correct metalation, including pre-assembling metal cofactors (which allows for easier distinction and delivery of specific metals), the use of metallochaperones (removing especially thermo-dynamically-favored elements like Cu / Zn from the accessible, intracellular trace metal pool), and the compartmentalization of trace metal metabolism (adjusting metal concentrations locally in order to direct binding to target proteins) (6, 10, 11).

*Chlamydomonas reinhardtii* is a unicellular green alga that has been widely used as a eukaryotic, photosynthetic reference system, and therefore exploited in our laboratory to study trace metal metabolism. It has a short generation time (∼ 6h), is a facultative heterotroph and can be grown to high densities (12). Chlamydomonas requires a broad spectrum of metal cofactors to sustain its photosynthetic, respiratory and metabolic capabilities, with Fe, Cu, Mn and Zn as the major second row metals involved in these processes. In the last 20 years, studies have revealed a repertoire of assimilatory and distributive transporters in Chlamydomonas, using biochemical and genomics approaches (13–17), discovered mechanisms for metal-sparing and recycling to ensure economy (18–21), and identified metal storage sites (2, 22–24).

Storage sites are crucial components of trace metal homeostasis. The capacity to sequester individual trace metals is important for controlling protein metalation, detoxification in situations of overload, and buffering during metabolic remodeling. Metal storage provides a selective advantage in competitive environments when transition metals are scarce. One well known storage site is ferritin, a soluble, mostly cytosolic (animals) or mostly plastidic (plants), 24-subunit oligomer that can oxidize ferrous to ferric Fe and store up to ∼4500 ferric ions in mineralized form in its core (25, 26). The importance of ferritin as an Fe store is well documented in eukaryotes. In Chlamydomonas, the pattern of expression of ferritin is more consistent with a role in buffering Fe during metabolic transitions, for example from phototrophy to heterotrophy during Fe starvation (23, 27). Vacuoles, lysosome-related and other acidic organelles, are equally important storage organelles in eukaryotes (28–33). In yeast and plants, these organelles can sequester metals for future use (34). Chlorophyte algae employ a set of smaller cytosolic vacuoles, including contractile vacuoles and acidocalcisomes. Contractile vacuoles (CV) manage the water content in the cytoplasm; in a fresh water alga that is predominantly facing hypotonic environments (35, 36) water is removed from the cell, potentially using potassium (K) and/or chloride (Cl) (37) to generate an osmotic gradient to attract water to the CV. Acidocalcisomes are lysosome-related organelles in the cytosol, defined by their low pH and high levels of calcium and polyphosphate (polyP) content (38, 39). In Chlamydomonas and other green alga this organelle may also contain K (40, 41). Acidocalcisomes can be identified as electron-dense granules by transmission electron microscopy (TEM), or by utilizing specific probes targeting either the low pH environment or the specific elemental makeup of the compartment (24, 42–44). In addition, acidocalcisomes have been visualized using their unique elemental signature by mass spectrometry-based (LA-ICP-MS or nanoSIMS) or X-ray fluorescence microscopy (XFM) (45, 46). These methods demonstrated that, at least in Chlamydomonas, the acidocalcisome can house high amounts of Cu and Mn in excess conditions (24, 47), making it a prime candidate for Fe storage as well.

XFM is a synchrotron-based technique that utilizes X-rays to produce photons in the object to be visualized. The energies of these fluorescent photons are element specific and can be used to determine elemental distribution and concentrations in whole cells or subcellular compartments (48–50). High energy X-rays (> 10 keV) penetrate biological material deep enough so that no sectioning is required. True elemental distributions are obtained when metabolic states are preserved rapidly using either vitrification or chemical fixation. XFM has previously been utilized to determine the elemental content of vacuoles of phagocytes infected with various Mycobacterium species (51), to demonstrate transient zinc relocation to the nucleus during macrophage differentiation (52) and the mobilization of Cu during angiogenesis (53).

In this work, we employed XFM with sub-100 nm spatial resolution to identify an Fe storage site in Chlamydomonas, determined the spatial distribution of multiple essential trace elements in chlorophyte algae, and quantified acidocalcisomal metal content in single cells in situations of Fe and Cu over-accumulation. We took advantage of a Chlamydomonas vacuolar transporter chaperone (*vtc1*) mutant (defective in polyP synthesis and hence also Ca content (43, 47)) to distinguish the role of polyP and Ca in acidocalcisome Fe sequestration. XFM also enabled the comparison of single-cell trace metal quotas to bulk quantification of Cu and Fe in corresponding cell cultures, allowing us to distinguish the nutritional state for Cu and Fe in individual cells.

## Results

### XFM is an excellent technique to visualize trace metal sequestration in microalgae

We first focused on Zn-deficiency in Chlamydomonas, where cells over-accumulate intracellular Cu, since this condition offers excellent signal to noise and the site of Cu(I) accumulation was already identified as the acidocalcisome (15, 24). ICP-MS/MS of 5×10^7^ Chlamydomonas cells collected from Zn-deficient cultures showed that their Zn content was reduced to ∼ 1/3 of the abundance compared to cells grown in Zn replete medium (Figure 1A). The Zn deficient cells had high intracellular Cu content, corresponding to ∼ 20 times the amount found in Zn replete conditions (Figure 1A), consistent with results from earlier studies in which a similar over-accumulation of Cu was observed when intracellular Zn levels dropped to < ∼1×10^7^ atoms/cell (15). Zn-deficient cells also accumulated Fe, to about 3 times the amount found in replete conditions, while the amount of intracellular P and Ca did not significantly change (Figure 1A). Previously, we showed that Cu is sequestered into distinct Ca-containing foci using a fluorescent probe to identify Cu(I) foci and complementary, direct elemental identification of Ca and Cu by nanoSIMS imaging of thin sections. Noting that elemental maps derived from nanoSIMS are quantified relative to another abundant element in the section, we sought a quantitative description of the organelle utilizing XFM.

**Figure 1:**
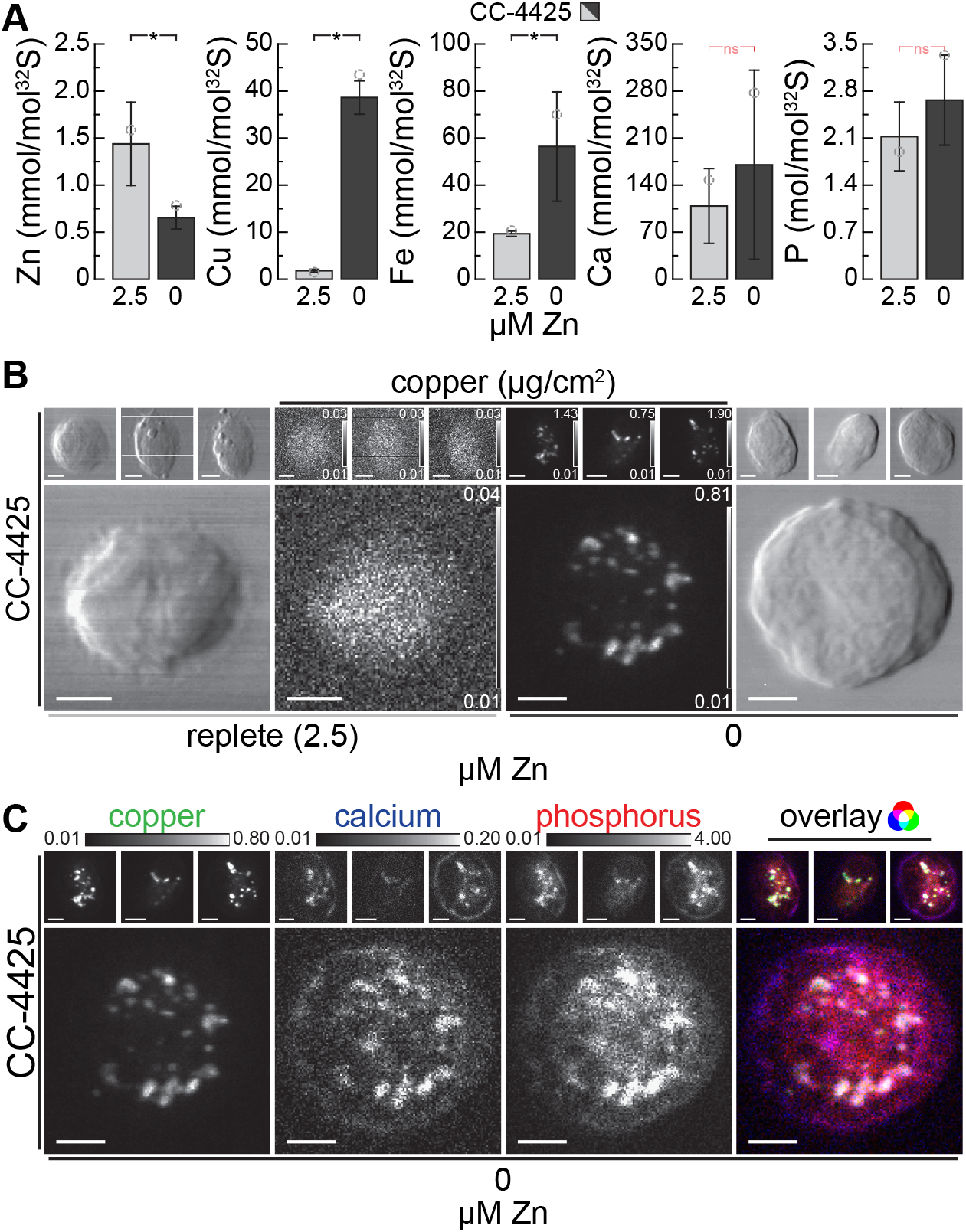
Zn deficient cells accumulate Copper (Cu), which is sequestered with Ca and P in acidocalcisomes. (A) Zn, Cu, Fe, Ca and P contents of *Chlam-ydomonas reinhardtii* wild-type strain CC-4425, as measured by ICP-MS/MS, either in the presence of 2.5 µM Zn (replete, light grey), or after two rounds of growth (5 generations each) in Zn deficient growth medium (Zn-deficient, dark grey). Elemental content is normalized to total cellular S content as a measure for total biomass, error bars indicate standard deviation of individual cultures (n≥3), the open circles indicate the elemental content of the specific batch that was used for X-ray fluorescence microscopy (XFM) analysis (Figure 1B, C). Asterisks indicate significant differences (t-test, P ≤ 0.05), non-significant (ns) outcomes are indicated in red. (B) Cu distribution in 4 individual CC-4425 wild-type cells, in Zn-replete (left) or Zn-deficient (right) conditions, measured by X-ray fluorescence microscopy at the Bionano-probe (APS). The images of chemically-fixed alga cells show cell outlines (outside panels) and Cu distribution (inside panels) ranging from the minimal (black) to maximal (white) Cu concentration in the individual image (in µg/cm^2^). The actual individual minimum and maximum concentration of Cu for each cell are denoted in the lower and upper right hand corner of each image, respectively, scale bars in the lower left corner indicate 2 µm. The cells were fixed and stored at room temperature, the images were acquired using fly-scan mode (continuous motion in the horizontal direction) at low temperatures with 70 nm step size and 300 ms dwell time per position. The fluorescence maps were created by performing peak area fitting for every position utilizing the MAPS software suite. (C) Cu, Ca and P distribution of the 4 wild-type cells in Zn-deficiency from (B), as well as the overlay of the 3 elements. All elemental distributions for all cells are depicted between shared, fixed minimal (black) and maximal (white) elemental concentrations, denoted above the images (in µg/cm^2^), white bars in the bottom left corner of each image indicate 2 µm. The rightmost image shows the overlay of the three elements, where Cu contributes the green, Ca the blue and P the red color.

Cu displayed different localization patterns in cells grown in Zn-replete vs Zn-deficient conditions. While Cu was distributed equally throughout the cell in replete conditions, it was found predominantly in foci in Zn deficiency (Figure 1B, Supplemental Figure 4A) similar to the earlier findings in TEM, nanoSIMS and fluorescent images. Ca, as described earlier (24), and also P were found to be co-localized with Cu to the foci, both in chemically-fixed and vitrified cells (Figure 1C, Supplemental Figure 4B). In general, vitrified samples had sharper intracellular structures compared to chemically-fixed material, which is especially pronounced for P and Ca. Both methods of preservation however preserved the Cu sequestration site and maintained the co-localization pattern sufficiently. Co-localization confirmed and further strengthens the conclusion that indeed acidocalcisomes, namely acidic, polyP- and calcium-rich, cytosolic vacuoles, were the storage site for Cu in Zn-deficiency. By confirming and extending our earlier findings, we conclude that XFM, using either chemically-fixed or vitrified algae, is a viable technique for analyzing intracellular metal storage compartments.

### Fe is sequestered in Chlamydomonas acidocalcisomes independently of polyP and Ca

Based on the data presented here and described previously in (24, 47), we recognize the acidocalcisome as a prime location for essential metal storage in Chlamydomonas. In addition to over-accumulation of Cu in Zn deficiency and Mn in excess Mn environments, we asked if the organelle might also store Fe. Fer1, the major isoform of ferritin in Chlamydomonas, was found to be less abundant in cells cultured in medium supplemented with excess Fe, indicating that other storage sites might primarily be used for Fe sequestration in Chlamydomonas (23). We used the following strategy to overload Chlamydomonas cells with Fe. First we subjected cells to a brief period of Fe-limitation (0.1-0.25 µM Fe in the medium), in which Fe assimilation components are dramatically up-regulated (17). Accordingly, upon supply of excess Fe (200 µM Fe, corresponding to 10-fold the amount supplied in a replete culture), the cells took up large amounts of Fe (Figure 2A, Supplemental Figure 1), resulting in transient over-accumulation of Fe to about 8-fold the amount found in typical replete cultures. Elevated Fe levels persist for about 48-72 h, before a lower, but still elevated state (two-to three-fold relative to a typical replete culture) is reached. The transiently over-elevated Fe levels span a period of 2-3 generations at peak abundance. During this time, Fe needs to be stored safely to avoid detrimental effects from Fe toxicity (8).

**Figure 2:**
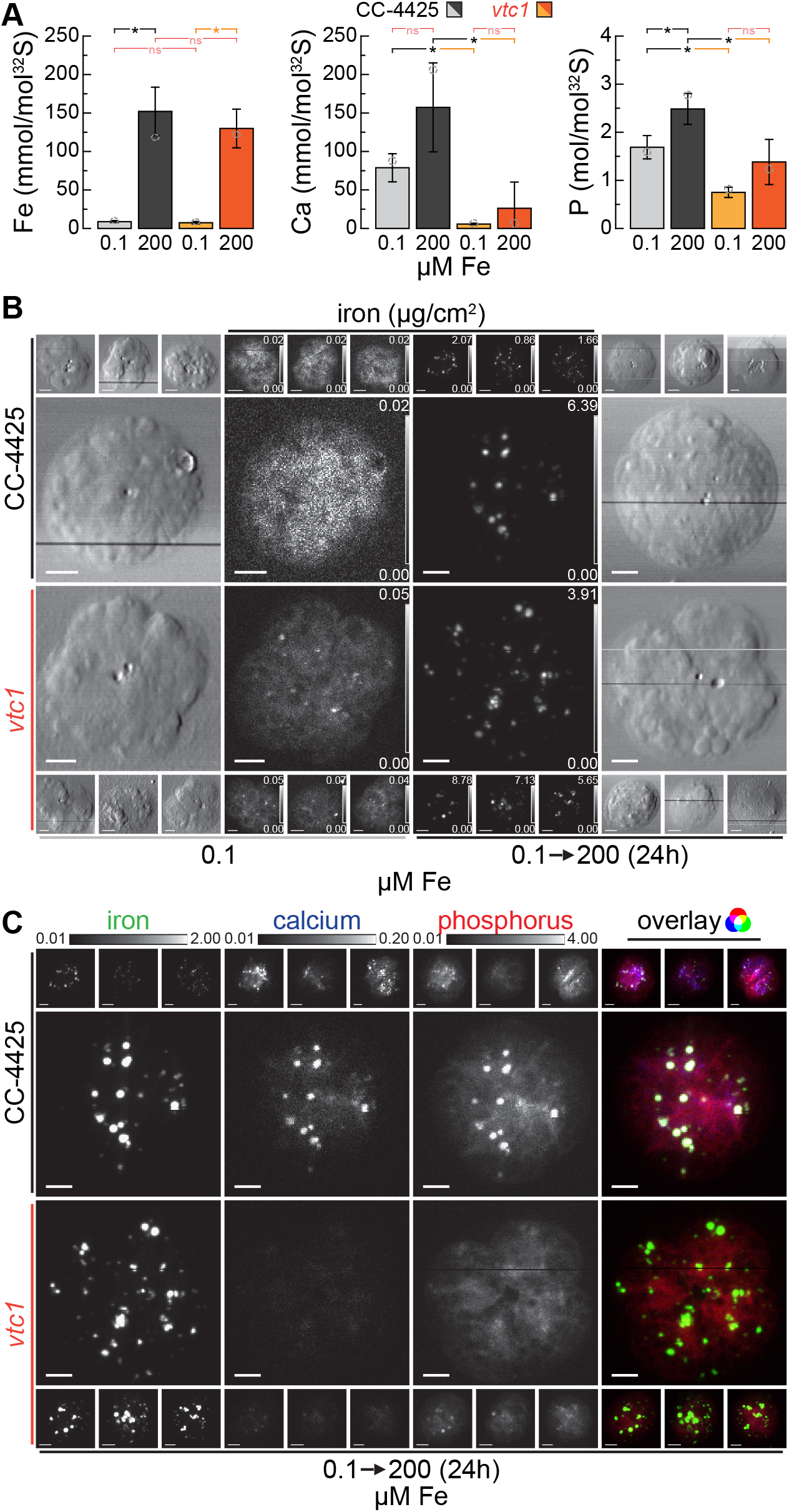
Acidocalcisomes serve as Fe storage sites during transient overload, acidocalcisomal Ca and polyP is not required for sequestration. (A) Total Fe, Ca and P contents of *Chlamydomonas reinhardtii* wild-type (CC-4425, grey bars) and the *vtc1* mutant strains (orange bars), as measured by ICP-MS/MS, either in Fe-limiting conditions (0.1 µM Fe) or 24 hours (2 doublings) after the addition of 200 µM Fe to Fe-limited cells (transient Fe over-accumulation). The elemental content is normalized to total cellular S content, as a measure for total biomass, error bars indicate standard deviation of individual cultures (n≥3), open circles indicate the elemental content of the specific batch used for X-ray fluorescence microscopy analysis in (B, C). Asterisks indicate significant differences (t-test, P ≤ 0.05), non-significant (ns) outcomes are indicated in red. (B) Fe distribution of 4 individual cells, in Fe-limited conditions or after 24 h of Fe– excess, measured by X-ray fluorescence microscopy at the Bionanoprobe (APS). Cells were prepared as described in Fig1B. The images of chemically-fixed alga cells show the cell outlines (outside panels) and the Fe distribution (inside panels) ranging from minimum (black) to maximum (white) Fe concentration in µg/cm^2^, actual minimum and maximum concentrations for each cell are noted in the lower and upper right hand corner of each image, white scale bars in the bottom left indicate 2 µm. (C) Fe, Ca and P distribution of the 8 (4 wild-type and 4 mutant) 24 h Fe–excess cells from Fig 2B, as well as overlay. All elemental distributions for all cells are depicted between shared, fixed minimal (black) and maximal (white) elemental concentrations, denoted above the images (in µg/cm^2^), white bars in the bottom left corner of each image indicate 2 µm. The rightmost image shows the overlay of the three elements, where Fe contributes the green, Ca the blue and P the red color.

In order to analyze the contribution of acidocalcisomes to Fe handling in Fe over-accumulating conditions directly, we analyzed the distribution of Fe in corresponding wild-type cells as well as in the *vtc1* mutant (43). The VTC complex is a multi-subunit membrane-spanning enzyme (54, 55) responsible for polyP synthesis in the acidic vacuoles (56, 57). In *Saccharomyces cerevisiae*, among the several proteins of the complex, only loss of VTC1 function results in a complete loss of the entire complex (55). Consistent with VTC1 function and earlier studies (47), Chlamydomonas *vtc1* mutants have reduced total cellular P, ∼ 40% relative to wild-type and the complemented strain (*VTC1-C2*), presumably attributed to reduced acidocalcisomal polyP content (Figure 2A, Supplemental Figure 1B). Interestingly, elemental analysis also showed a dramatic reduction of cellular Ca in the mutant, compared to both the wild-type and complemented strain (Figure 2A, Supplemental Figure 1B), suggesting that most of the cellular Ca is associated with polyP in acidocalcisomes. Wild-type, *vtc1* mutant and complemented strain (*VTC1-C2*) showed similar intracellular Fe accumulation (∼ 8 times the amount in Fe-replete conditions) after excess Fe addition to Fe-starved cells (Figure 2A, Supplemental Figure 1A/B). Furthermore, temporal and concentration-dependent elemental profiles indicated similar behavior of wild-type, mutant and complemented strains (Supplemental Figure 1) upon Fe resupply. In both, mutant and wild-type cells, Fe is concentrated into distinct foci in the Fe overload situation, while at low intracellular Fe levels, the Fe is more evenly distributed throughout the cell (Figure 2B), presumably largely in Fe-binding proteins. According to XFM, Fe foci contained also both Ca and P in wild-type cells, suggesting the acidocalcisome as the likely site of Fe sequestration (Figure 2C, Supplemental Figure 5). In the *vtc1* mutant, where acidocalcisomes lack polyP and Ca, the same sequestration pattern was observed (Figure 2C, Supplemental Figure 5). Additionally, in wild-type algae but not in the *vtc1* mutant, P accumulates alongside Fe (Figure 2A).

As mentioned above, another situation in which Fe over-accumulates in cells is under Zn starvation, where the Fe content is comparable to cells exposed to long-term Fe excess, ∼ two to three times the quota of typical Zn and Fe-replete cultures (Figure 1A and 3A). When we tested Zn deficient wild-type and *vtc1* cells for the site of Fe accumulation, we found the same pattern as in the Fe overload situation. Specifically, a punctate pattern of Fe co-localized with Ca and P in wild-type cells, and no impact of the *vtc1* mutant on the Fe content or pattern of distribution (Figure 3A/B). On the other hand, the *VTC1* genotype did affect Mn over-accumulation in Zn deficiency (Figure 3B) and upon Fe overload (Supplemental Figure 1B), just as it did in situations of Mn overload (47). Like Fe, Cu accumulation in Zn deficiency was also not impacted by loss of acidocalcisomal polyP synthesis (Figure 3A). In all experiments, Ca and P were reduced in the *vtc1* mutant but restored in the complemented strains, confirming that Ca accumulation requires *VTC1* function (Figure 2A and 3A, Supplemental Figure 1A/B). Taken together, we conclude that over-accumulated Fe, independent of the absolute amount (two to 8-fold), is housed in the acidocalcisome, defined by its Ca and P contents. We further conclude that acidocalcisomal polyP and Ca are not required for Fe accumulation in the organelle, in contrast to Mg and Mn accumulation (47, 58).

**Figure 3:**
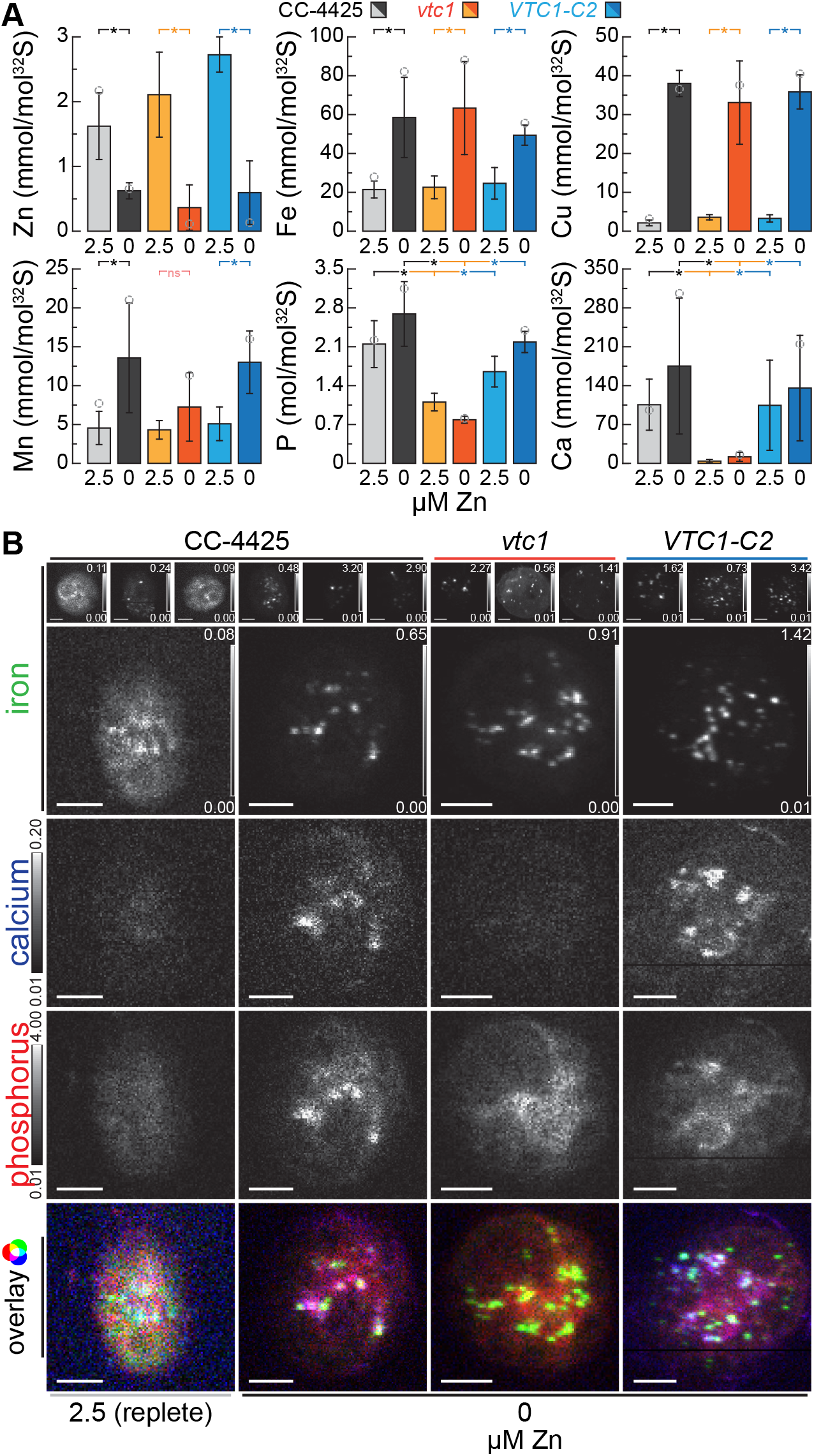
Fe accumulated in Zn deficiency is sequestered in acidocalcisomes, independent of Ca and polyP. (A) Total Zn, Fe, Cu, Mn, Ca and P contents of *Chlamydomonas reinhardtii* wild-type (CC-4425, grey bars), the *vtc1* mutant (orange bars) and VTC1-C2 complemented strain (blue bars), as measured by ICP-MS/MS, either in the presence of 2.5 µM Zn (replete, lighter colors), or after two rounds of growth (5 generations each) in Zn deficient growth medium (Zn-deficient, darker colors). The elemental content is normalized to total cellular S content, as a measure for total biomass, error bars indicate standard deviation of individual cultures (n≥3), open circles indicate the elemental content of the specific batch used for X-ray fluorescence microscopy analysis (Figure 3B). Asterisks indicate significant differences (t-test, P ≤ 0.05), non-significant (ns) outcomes are indicated in red. (B) Fe distribution in 4 individual cells from the wild-type (CC-4425), the *vtc1* mutant and VTC1-C2 complemented strain (blue bars) in Zn-replete or Zn-deficient conditions, as measured by X-ray fluorescence microscopy at the Bionanoprobe (APS). Cells were prepared as described in Fig1B. The minimum (black) to maximum (white) Fe concentration in µg/cm^2^ is depicted in the lower and upper right hand corner of each image, elemental distributions for Ca and P in all cells are depicted between shared, fixed minimal (black) and maximal (white) elemental concentrations, denoted left of the images (in µg/cm^2^), white scale bars in the bottom left indicate 2 µm. The bottom image shows the overlay of the three elements, where Fe contributes the green, Ca the blue and P the red color.

### Cu and Fe quota of single cells can be determined accurately using XFM

Because XFM is element-specific and quantitative, we sought to compare intracellular Cu and Fe quotas at single-cell resolution with bulk ICP-MS/MS measurements of 5×10^7^Chlamydomonas cells cultured in the same conditions. Our experimental conditions offer a dynamic range of two orders of magnitude with respect to Cu and Fe contents. In multiple XFM analyses, 5-8 individual cells were each analyzed in Fe-limited (0.1 µM Fe), Fe-replete (20 µM Fe), Fe-accumulating (Zn deficient, 20 µM Fe, 0 µM Zn) and Fe-over-accumulating (0.1 µM Fe, 200 µM Fe addition) conditions. Data from four experimental runs, over a period of two years, are presented here. With the exception of Fe-replete conditions, all growth conditions were measured multiple times from independent experiments or different strains, either rapidly vitrified or chemically fixed. This is reflected by individual data points in the graphs (Figure 4A/B). The total intracellular Fe content in the individual cells, as measured by XFM, correlated well with the intracellular content of the bulk-cultures, as measured with ICP-MS/MS (Figure 4A). Quantitative XFM measurements allowed accurate measurement of single-cell Fe content, and therefore is indicative of the Fe-nutritional state of an individual cell. For Cu we analyzed conditions representing three different stages of Cu accumulation: Cu-replete (2 µM Cu), Cu-accumulating (2 µM Cu, 0.1 µM Fe before and after 200 µM Fe addition) and Cu over-accumulating conditions (2 µM Cu, 0 µM Zn). Similar to Fe, we noted a remarkable correlation (r^2^ > 0.8) between single-cell and bulk measurements (Figure 4B). Again, XFM on its own allows the distinction between various states of Cu nutrition and accurately assesses the total intracellular content. Quantitative data from both, chemically-fixed (solid symbols and errors, Figure 4A/B) and vitrified (dashed symbols and errors, Figure 4A/B) samples, were obtained and compared equally well with ICP MS/MS data, both for Fe and Cu content. Synchrotron beamline time allocation is a crucial limiting factor with respect to the number of cells that can be completely scanned at the desired spatial resolution. Similar to other imaging-based techniques, we made an effort to “randomly” choose cells of various sizes for high resolution imaging, insofar as this was possible with the small number of cells that was analyzed per condition. The size range reflects the size variation in batch cultures from an asynchronous population. This allows us to control for cell-cycle-stage specific dynamics, but it can also contribute to the variance when elemental abundance data is normalized per cell, like in this case. For other elements we did not actively seek to manipulate their intracellular accumulation (Ca and P levels were altered in *vtc1* mutant strains as described above, Mn was accumulating in Zn deficiency to two to three times the amount of replete levels), and therefore their intracellular abundance did not span multiple orders of magnitude. In general P, S and Ca abundance in individual cells, as measured by XFM, matches the levels measured by ICP-MS/MS. For Mn, the abundance measured by XFM was low, potentially from interference, but the relative content correlates with increasing amounts of intracellular Mn (Supplemental Figure 6). For Zn, background levels were too high to allow for a quantitative assessment (Figure 4C). Taken together, we conclude that Fe, P, S, Ca and Cu levels of individual cells can be accurately quantified using XFM and can be used to predict the nutritional status of single cells in environmental samples. Mn and Zn measurements allowed only for the analysis of spatial distribution and relative changes using the present setup.

**Figure 4:**
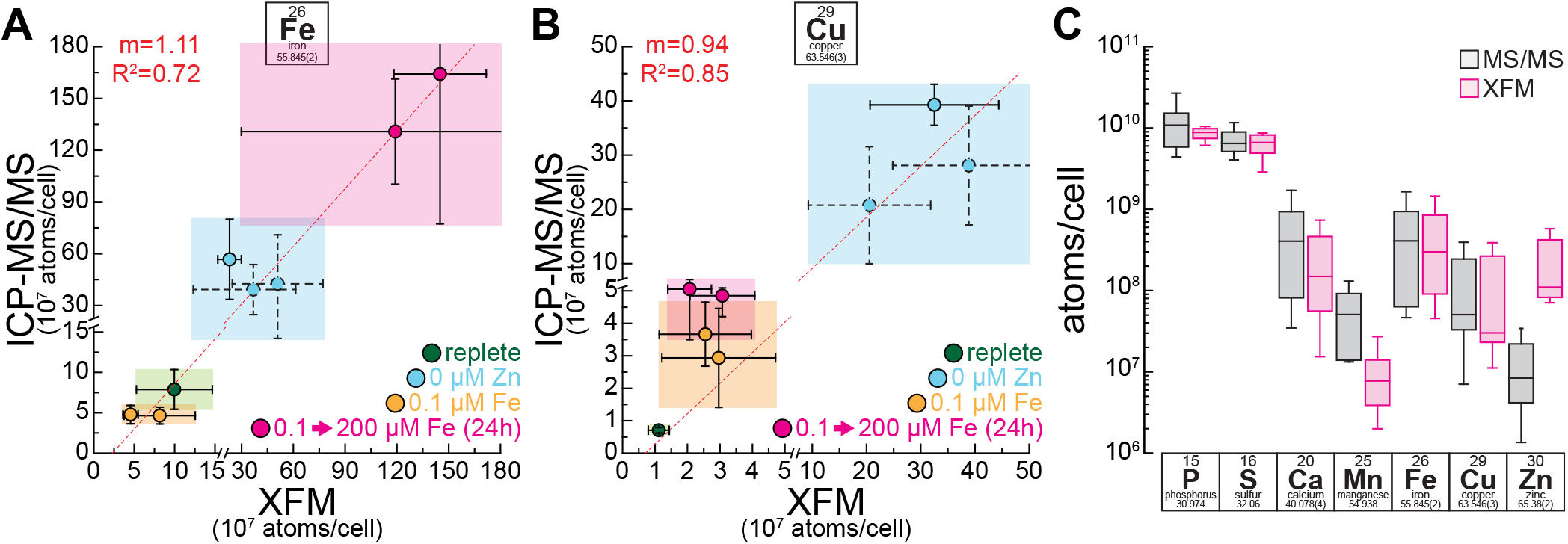
Single cell Fe and Cu content accurately recapitulates batch culture data. (A/B) Correlation of Fe (A) and Cu (B) content as measured by X-ray fluorescence microscopy (x-axis) and ICP-MS/ MS (y-axis). Error bars in x and y direction indicate standard deviation in the measurements between at least 4 individual cells (XFM) or between at least 3 independent cultures (ICP-MS/MS). Green areas indicate cells/cultures from replete conditions, blue colored areas indicate Zn-deficient conditions (Cu over-accumulation, Fe two to three times replete levels), pink areas indicate 24h of Fe-excess (Fe over-accumulation, Cu two to four times replete levels) and orange areas indicate Fe-limited conditions (low Fe, Cu two to four times of replete levels). Solid outlines of data points and error bars are associated with chemically fixed samples, dashed outlines with samples that were vitrified. Both axes were broken at the same position to magnify samples with lower intracellular Fe or Cu levels. A linear regression analysis was performed in OriginPro (v9.1), the resulting regression line is presented in the image as a red dashed line, interrupted only from the axis break. R^2^ and slope (m) of the analysis are denoted in red in the top left hand corner. (C) Box-plots for elemental content of different intracellular elements, as measured by ICP-MS/MS in between all experimental conditions (grey) and in between all imaged cells in all experimental conditions (pink). From bottom to top, horizontal lines indicate 5%, 25%, 50%, 75% and 95% quantile.

### S performs better than P in facilitating comparisons between bulk cells and elemental distributions

In order to facilitate the comparison between trace metal abundance data acquired by ICP-MS/MS and the spatially distributed XFM data, we explored the possibility of using phosphorus (P) or sulfur (S) for normalization in addition to cell numbers (for bulk elemental content) and cell area (for spatially distributed elemental maps). Both P and S were promising candidates for normalization because both elements’ distribution can be accurately analyzed in XFM (59, 60) and accurately quantified with ICP-MS/MS in mass-shift mode (61, 62). Using cellular P or S content for normalization allows us to correct for synchronization artifacts, introduced from experimental perturbations, for example in experiments where high amounts of Fe were added back to Fe-starved cells (Supplemental Figure 1A, 6-h time point). Additionally, P or S normalization allows us to account for differences in cell size, resulting from random selection of cells of a range of different sizes during XFM analysis. Elemental distribution maps for S or P might also facilitate the analysis of enrichment of elements within storage compartments, when equally distributed they could better reflect depth of the cell at each position of the image, compared to using the area alone (Supplemental Figure1/2, also (52)).

First, we analyzed the amount of intracellular S and found that it was not affected by the perturbations used in this work (Supplemental Figure 1A, B). The total P content in some of the strains used in this study (*vtc1*) was reduced, but P was not affected by the Fe regime (Figure 2A, Supplemental Figure 1B) or by Zn status (Figure 3A). Consequently, in a single genotype, changes to trace element contents between conditions were not affected by the choice of normalization (P, S or cell number (see Supplemental Figure 1A). When comparing to biomass, measured as non-purgeable organic carbon content (NPOC, Supplemental Figure 2), S content correlated better with the carbon content of cells than did P. S performed equally well in both genotypes for the correlation with biomass (Supplemental Figure 2A), while for P the correlation improved markedly when *vtc1* was omitted (Supplemental Figure 2B), although it did not reach to the level of correlation observed for S. We noted also that P (Figure 1C, Figure 2C, Figure 3B) was less evenly distributed than S in cells (Supplemental Figure 3), in part because of its accumulation in the acidocalcisomes, but also because of segregation in other organelles, including the chloroplast, which appears to have less P compared to other compartments. Taken together, P content varies between strains, exhibits a more complex intracellular distribution and less accurately captures biomass. We conclude that S is a better choice for area-dependent normalization. P does generally correlate well with biomass, leaving the possibility of its use for normalization in a different context.

### Two types of acidocalcisomes can be distinguished based on their elemental composition

In order to identify the elemental composition of the compartments that contain over-accumulating trace metals, we used a k-means cluster analysis feature in the MAPS software (63) to identify continuous, secluded areas with high Cu or Fe contents in various metal-accumulating conditions. In Zn-deficient cells, as much as 70% (∼ 55 % on average) of the intracellular Cu was found in areas containing storage compartments. We found Fe, Ca, P and Mn to be enriched in these areas, either relative to the cell area or the S content present in these areas (Figure 5A). In Fe over-accumulating conditions up to 80% (∼ 65 % on average) of the total cellular Fe was found in areas containing storage compartments. Again, we found Ca, P and Mn, but Zn instead of Cu to co-localize with Fe. The concentrations of P and Ca within these areas was comparable between Cu- and Fe-accumulating conditions, while the combined concentration of Fe and Cu in the areas containing storage compartments was ∼ three-fold higher in Fe accumulating conditions (Figure 5B). In addition, the metal composition of these areas was significantly different in Cu- and Fe-accumulating conditions. In Zn deficiency, the areas contained mixed trace metal populations of predominantly Fe and Cu, while Fe alone was the dominating metal in Fe over-accumulating conditions. When analyzing intracellular trace metal abundance outside the metal storage sites (Figure 5C), similar Fe concentrations were maintained between replete, Fe-over-accumulating and Zn-deficient conditions, indicating that excess Fe is successfully sequestered in the acidocalcisomes. Intracellular Fe levels were however reduced ∼ 5-fold in Fe-limited conditions (Figure 5C), indicative of the operation of Fe sparing responses (19). For Cu on the other hand, in Zn-deficient (copper-accumulating) conditions, intracellular Cu levels outside the acidic vacuoles are also elevated, ∼10-fold, consistent with an initiating event for Cu accumulation at a cytoplasmic site.

**Figure 5:**
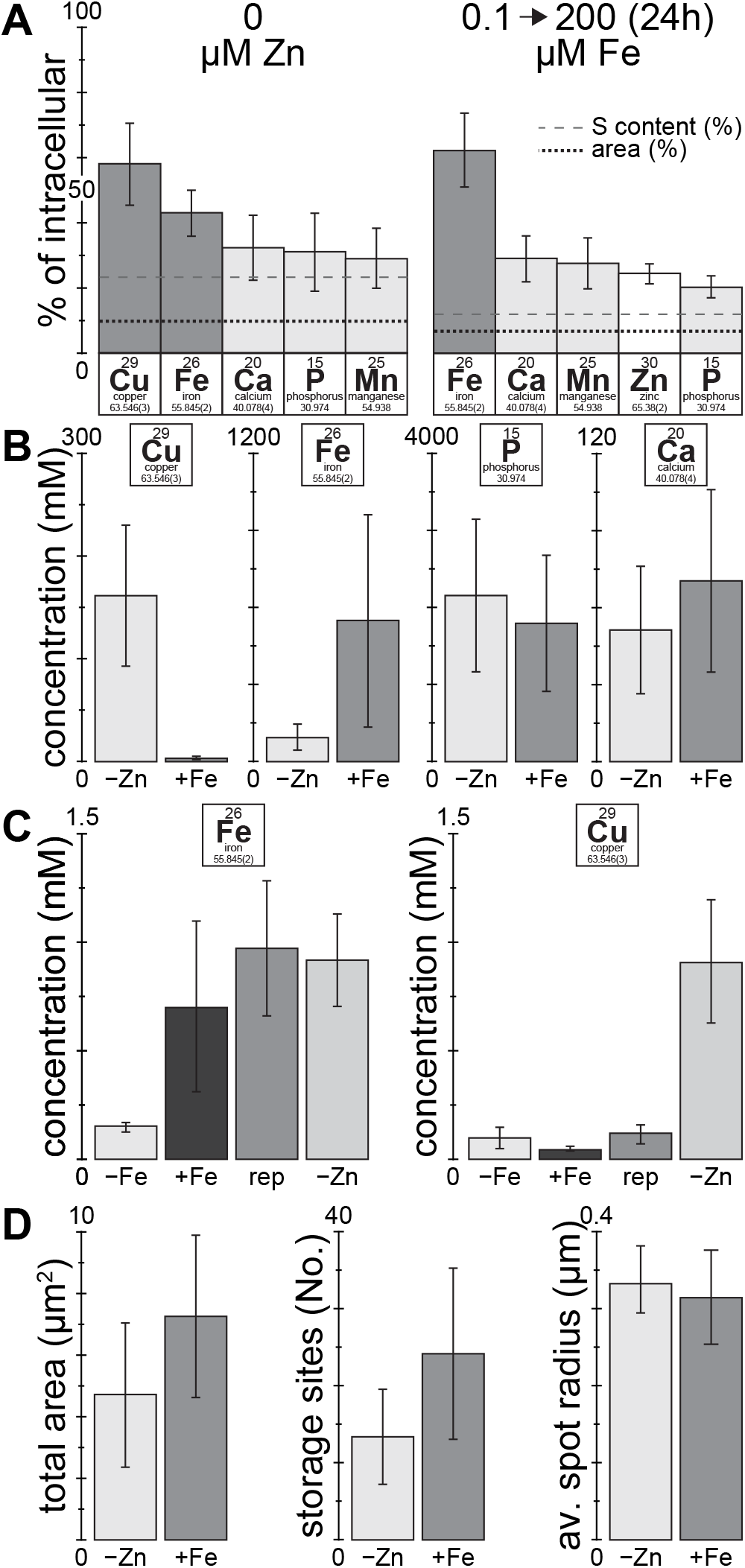
Most of the Fe and Cu overload is in the acidocalcisome. (A) Fraction of total cellular elemental content enriched within areas of high Cu (in Zn deficiency) or high Fe content (in 24 h of Fe-excess), respectively. Areas containing storage compartments were determined by a *k*-means clustering approach identifying areas of continuous, high Cu or Fe concentration, respectively, using the MAPS software. The black, dotted line indicates the fraction of the cell area containing storage compartments, the grey, dashed line indicates the fraction of the S content (as a proxy for the cell volume) within the areas containing storage compartments. Included are only elements enriched in the areas containing storage compartments. Error bars indicate standard deviation from at least 4 individual cells. (B) Average Cu, Fe, Ca and P concentration within the acidocalcisomes in Zn deficiency (Cu-over-accumulation) or Fe-over-accumulating conditions. For calculation of the elemental concentration within the acidocalcisome, the volume of the organelle was extrapolated from the average dimensions obtained from the areas containing storage compartments, and the organelles were assumed to be spherical. Error bars indicate standard deviation between organelles from at least 4 individual cells. (C) Average intracellular Fe and Cu concentration outside the storage regions, in replete, Fe-limited, Fe over-accumulating and Zn-deficient conditions. Error bars indicate standard deviation between at least 4 individual cells. (D) Total intracellular area containing storage compartments, the number of individual regions of continuous, high Cu or Fe concentration, and their extrapolated radius, derived from the total area covered and the number of regions within a cell. Error bars indicate standard deviation between at least 4 individual cells.

Acidocalcisomes covered ∼50% more cellular area in Fe over-accumulating conditions, compared to Cu accumulating conditions (Zn-deficiency) (Figure 5D). In both conditions despite more such organelles in Fe accumulating conditions, the average acidocalcisome had a similar radius of ∼ 0.3 µm (Figure 5D). In summary, the size (diameter), Ca and P concentration of acidocalcisomes were retained between the two physiologically-distinct metal accumulating conditions, whereas the number of acidocalcisomes and the abundance and composition of the accumulated material were altered, allowing the cell to accommodate different demands for trace metal sequestration.

## Discussion

The acidocalcisome is an acidic vacuole in the cytosol of most eukaryotic organisms, high in polyP and Ca content (42). In Chlamydomonas, the organelle is more prevalent when cells reach stationary phase (44) and in situations of nutrient limitation, for example in S deficiency (43), N deficiency (44) and Zn deficiency (24). All of these perturbations are associated with a strong reduction in the growth rate, resulting from the absence of a single, essential nutrient, therefore presenting the opportunity for cells to accumulate other, non-limiting nutrients in excess amounts. While the role of the acidocalcisome for nutrient storage in N and S deficiency is still unclear, a role in trace metal homeostasis is established. Specifically, it has been demonstrated that Zn-deficient Chlamydomonas acidocalcisomes facilitate trace metal storage and harbor excess amounts of other non-limiting trace elements, most prominently Cu, but also Fe and Mn (Figure 1 and 3). Nutrients accumulated in these situations are accessible, at least in the case of Cu, which can be later utilized to facilitate growth when Cu becomes limiting (64). More recently, the acidocalcisome was found to store excess Mn, which can also be utilized later (47) and to store excess Fe, when indulged with Fe after a brief period of Fe limitation (Figure 2). It is noteworthy that the Fe and Mn-accumulating conditions are distinct from those used in previous work. In contrast to the Cu over-accumulating situation, the growth rate of the alga is not limited during Fe and Mn accumulation. For Fe in fact, the growth of the cells, which was initially Fe-limited, improved dramatically by adding excess amounts of Fe. For Mn, a subtle but significant increase in growth rate was observed with higher Mn concentrations (Tsednee *et al*. 2019, Figure 1A). The increase in intracellular Fe and Mn contents was solely driven by an increase in external (environmental) Fe or Mn. We suggest that the role of the acidocalcisome in these conditions is to sequester intracellularly un-needed (and hence excess) trace metals in order to maintain healthy Fe and Mn homeostasis in the cytoplasm.

We were able to quantitatively trace all intracellular metal pools using a combination of XFM and ICP-MS/ MS. Vitrified and chemically-fixed alga cells showed equally well correlation with bulk data. We found that up to 80% of total cellular Fe is stored in acidocalcisomes when cells are over-accumulating Fe. This means that almost all of the excess Fe, the amount that exceeds the native, replete Fe content in cells, is sequestered within the acidic vacuoles. The same is true in Zn deficient conditions, where cells accumulate two to three times more Fe, and consequently up to 60 % of total Fe can be found in acidocalcisomes, again reflecting the excess part of the Fe quota. This was confirmed by a quantitative analysis of the Fe concentrations inside the cells, but not sequestered within the vacuoles, where both Fe accumulating conditions showed replete levels of Fe within the cell. This only leaves a limited role for other Fe storage sites in those conditions, for example the two ferritins Fer1 and Fer2 in Chlamydomonas. The abundance of the predominant plastid ferritin (Fer1) was shown to be reduced in high Fe conditions (23, 65), more consistent with a role of ferritin in buffering Fe released form intracellular sources for redistribution when Fe-limited (65). Chlamydomonas Fer2, which is also localized to the chloroplast and ∼ 70-fold less abundant, was not reduced in abundance when Fe is more abundant in the medium, but also did not accumulate (23). Given the Fe distribution data (Figure 2B), we did not note any accumulation of Fe in the chloroplast, leading us to conclude that Fer1 and 2 only have a minor role in Fe sequestration upon over-accumulation, if any. Many organisms in the Viridiplantae, which include chlorophyte algae and plants, harbor small gene families of ferritin, allowing for tissue-specific expression and regulation in response to different stimuli (66). In contrast to Chlamydomonas, ferritins in Arabidopsis and maize, for example, are induced upon exposure to elevated Fe and they do contribute to Fe storage (67, 68).

For Cu, the situation for storage is different. Lysosome-related organelles play a crucial role in Cu sequestration in eukaryotes, from yeast to mammals (69, 70). For example, storage and removal of excess Cu from liver hepatocytes is mediated via lysosomes (71); there Cu is associated with S, from metallothionein (72). Similar Cu concentrations (> 80 mM) in areas of high Cu concentration have been observed in tumorous rat livers (73), although these areas of high Cu concentration were vastly bigger in size (> 10 µm, compared to the < 1 µm acidocalcisomes in Chlamydomonas). Additionally, in higher eukaryotes, Pushkar et al. identified Cu storage vesicles (CSV) in astrocytes within the periventricular region of the lateral ventricle in rat and mouse brains (74). Cu concentrations within this compartment were also in a similar range to that observed here (100-300 mM). However, similar to liver cells, the sequestered Cu in astrocytes showed co-localization with S, probably from metallothionein, rather than with Ca or P. In Chlamydomonas, in both Zn deficient and Fe-limited conditions, total Cu levels were elevated. The majority of Cu in Zn deficient conditions is localized to acidocalcisomes (∼55 % on average, up to 70% in individual cells, Figure 5), but that does not capture all of the excess intracellular Cu. Our findings suggest that at least 30% of the accumulating Cu in Zn deficiency is bound by different means, either proteins or small ligands. Additionally, in Fe-limited conditions, Chlamydomonas cells increase their Cu quota. This increase in Cu is of similar extent compared to the increase of Fe in Zn deficient conditions (both increase two-to four-fold). While the increase in Fe can fully be explained by vacuolar Fe storage, for Cu, sequestration of the additional Cu in acidocalcisomes was not observed (Figure 4A), suggesting other explanations for the higher cellular Cu quota. One possibility is that in Fe-limited cells Cu accumulates to serve physiological needs; for instance to accommodate elevated amounts of FOX1, a periplasmic multicopper oxidase involved in high-affinity Fe uptake. With 6 Cu atoms and dramatic up-regulation in Fe-limited cells, FOX1 can be a substantial (∼ 12% of the quota) Cu-housing sink in the cell (14,18). Zn-deficient Chlamydomonas cells where *FOX1* expression also increases ∼10-fold also have increased intracellular Cu outside the acidocalcisomes (Figure 5C).

In summary, we utilized XFM to identify the acidocalcisomes as the storage site for Fe and Cu in situations of trace metal over-accumulation. We also quantified the elemental contents of whole cells and acidocalcisomes, and are therefore able to distinguish the nutritional state of an alga at the level of a single cell. This is promising not only for trace metal distribution studies, but allows for comparative single cell work, since the technique is non-destructive. Combined with upcoming upgrades to XFM capacity, increasing either throughput or resolution, studies comparing the single cell nutritional status with molecular indicators will be feasible.

## Methods

### Materials, strains and culture conditions

*Chlamydomonas reinhardtii* strains CC-4533 (CMJ030), CC-4425 (D66+), CC-5321 (*vtc1-1*, 43), CC-5324 (*VTC1-C2*, complemented *vtc1-1*, 43) were used for the experiments in this study. Cultures were cultivated in replete or trace metal-deficient medium, as indicated, in 250 mL Erlenmeyer flasks containing 100 mL TAP medium with a revised trace element solution, as described previously (77, 78). The cultures were grown with constant agitation in an Innova 44R incubator (160 rpm, New Brunswick Scientific, Edison, NJ) at 24°C in continuous light (90 µmol m^−2^ s^−1^), provided by cool white fluorescent bulbs (4100 K) and warm white fluorescent bulbs (3000 K) in the ratio of 2:1. All nutrient deficient media were prepared in flasks that were treated with 6N hydrochloric acid (analytical grade) for at least 12 h, to reduce the trace metal background from the vessel and ensure accurate metal contents across replicates dependent on the addition of trace metals and not impurities. The acid-washed flasks were rinsed 6 times with Milli-Q water before use to remove traces of the acid. Single-use plastic serological pipettes and spatulas and similarly acid-washed graded cylinders were exclusively used during the preparation of any of the solutions in this study. Cu over-accumulation in Zn deficiency was achieved by subjecting replete Chlamydomonas cultures to two consecutive rounds of dilution to an initial concentration of 1×10^5^ cells/ml, both times in medium that was prepared without the addition of Zn. Fe over-accumulation was achieved by diluting replete-grown, mid-log Chlamydomonas cultures once to 1×10^4^ cells/ml in 0.1 µM Fe media, before increasing the concentration in the culture to 200 µM Fe when the culture reached a density of 1×10^6^ cells/ml. The cell density reached between 3-6×10^6^ cells/ml 24 h after the addition of Fe.

### Total non-purgeable carbon and total nitrogen content analysis

Total, non-purgeable organic carbon content of the cells was determined as described in (79) with minor modifications on a Shimadzu TOC-L/TN CSH instrument. In brief, 3×10^7^cells were collected by centrifugation at 3100 *g* for 3 min, briefly washed once in 10 mM Na-phosphate (pH 7.0) and then resuspended in 900 µL of 3 M HCl (3.33×10^7^ cells/mL). Cells in HCl were incubated for 16 h at 65°C with constant agitation before being subjected to non-purgeable organic C and total N analysis. The acidified samples were diluted to 3×10^5^ cells/mL with Milli-Q water before being cleared of inorganic C by sparging with purified air and the determination of the non-purgeable organic C (NPOC) and total nitrogen (TN) content. Each sample was measured at least 3 times with a nondispersive infrared gas analyzer (CO_2_) and a chemiluminescence gas analyzer (TN). The peak area was calculated and compared with a standard curve from 0.5 to 25 ppm C (from potassium hydrogen phthalate) or N (from potassium nitrate) using the TOC-Control L software version 1.0 (Shimadzu). Measured concentrations did not exceed the calibration range and did not exceed 5% relative standard deviation between the individual technical replicates.

### Quantitative trace metal, S and P composition analysis

The trace metal, S and P composition was determined by ICP-MS/MS as described in (80) with minor modifications. In brief, 5×10^7^ Chlamydomonas cells of a culture during logarithmic growth phase (at a cell density between 1-5×10^6^ cells/mL) were collected by centrifugation at 3100 *g* for 3 min in a 50-mL Falcon tube. The cells were washed three times in 1 mM Na_2_-EDTA, pH 8 (to remove cell surface-associated metals), and once in Milli-Q water (to remove the EDTA) and transferred into a 15-mL Falcon tube in the process. The cell pellet, after removing the water, was overlaid with 143 µL of 70% nitric acid (trace metal grade, A467-500, Fisher Scientific) and incubated first at room temperature for 24 h and then at 65 °C for 4 h before being diluted to a final nitric acid concentration of 2 % (v/v) with Milli-Q water. Aliquots of fresh or spent culture medium were treated with nitric acid to a final concentration of 2 % (v/v). Metal, S and P contents were determined on an Agilent 8800 Triple Quadrupole ICP-MS/MS instrument, in comparison to an environmental calibration standard (Agilent 5183-4688), a S (Inorganic Ventures CGS1) and P (Inorganic Ventures CGP1) standard, using ^89^Y as an internal standard (Inorganic Ventures MSY-100PPM). The levels of all analytes were determined in MS/MS mode, where ^39^K, ^40^Ca, ^56^Fe and ^78^Se were directly determined using H_2_ as a cell gas, ^23^Na, ^24^Mg, ^55^Mn, ^59^Co, ^60^Ni, ^63^Cu, ^66^Zn and ^95^Mo where directly measured using He as a cell gas, while ^31^P and ^32^S were determined via mass-shift from 31 to 47 and 32 to 48, respectively, utilizing O_2_ in the collision / reaction cell. The average of 4-5 technical replicate measurements was used for each individual biological sample. The average variation in between the technical replicate measurements was below 2 % for all individual experiments and never exceeded 5 % for an individual sample.

### X-ray Fluorescence Microscopy (XFM)

3×10^6^ Chlamydomonas cells of a culture during logarithmic growth phase (at a cell density between 1-5×10^6^ cells/mL) were collected by centrifugation at 16,100 *g* at room temperature for 15 sec in a 1.5-mL Eppendorf tube. The cell pellet was washed twice in 1 x phosphate-buffered saline (PBS) buffer (137 mM NaCl, 2.7 mM KCl, 10 mM Na_2_HPO_4_, 1.8 mM KH_2_PO_4_) and resuspended either in 0.3 mL Milli-Q water for vitrification or 4 % para-formaldehyde (EM grade #15710 from EMS) in 1x PBS for chemical fixation. 100 µL of the cell suspension was then applied to a poly-L-Lysine coated silicon nitride membrane window (5 x 5 x 0.2 mm frame, 2 x 2 x 0.0005 mm Si_3_N_4_ membrane, Silson) and allowed to settle for 30 min at room temperature. After the supernatant was removed by gentle suction, the window was either directly mounted into an FEI Vitrobot Mark IV freezer for vitrification in liquid ethane (blot time 3 sec, blot force 2, blot total 1, wait and drain time 0 sec) or washed twice with 1x PBS, fresh 0.1 M Ammonium Acetate and Milli-Q water. Windows with chemically fixed cells were air dried and stored at room temperature, windows with vitrified cells were kept in liquid nitrogen storage until measurement at low temperatures. The data presented here are from four individual sessions (48-72 h measurement time each) at the Bionanoprobe (46), over the period of two years, during which the analyzer was localized either at beamline 21-ID-D or beamline 9-ID-B at the Advanced Photon Source (APS) of the Argonne National Laboratory. Direct excitation of atomic K transitions was aimed for by tuning the incident x-ray energy to 10 keV, which allows efficient excitation of elements in the elemental table up to Zn (z=30). Individual, large Chlamydomonas cells can be up to 15 µm in diameter before division, which is well below the 50–100 µm that can be penetrated by XFM with high spatial resolution (51). Whole cells where therefore analyzed without sectioning. Two rounds of coarse scans were performed to identify cell locations on the SiN windows, with a pixel size between 0.5-2 µm in x and y direction. Then, a high resolution scan of the individual cells was initiated for all images; a spatial resolution between 60 and 80 nm with a dwell time of 200 – 300 ms per pixel was used. With an average scan area of 10 x 10 µm, this results in measurement times up to 3 h per cell for each high resolution. By normalizing the recorded fluorescent spectrum with data obtained from a calibration standard, fully quantitative maps of the elemental distribution are obtained for each of the individual elements.

### Data analysis and software

X-ray fluorescence data were fitted and analyzed using the MAPS software package (63). Exported images were further edited and assembled in Adobe Photoshop and Illustrator.

For the quantitative analysis we defined three areas in the images, “whole cells”, which were further divided into “areas containing storage compartments” and “intracellular areas outside of storage regions”. Areas containing storage compartments, in all cases, were contained within the boundaries of the cells. The borders for the whole cells were identified in the images in the outside panels (Figure 1–3), and were manually replicated in the MAPS software in order to determine the amount of all the material contained within the cells and the area that the cell covers. These data were used for comparisons with ICP MS/MS data (Figure 4) of elemental content and contributed to the quantitative description of the metal storage compartment (Figure 5). “Areas containing storage compartments” were determined automatically, as continuous areas of high Cu concentration in Zn deficiency or Fe concentration during Fe accumulation, respectively, using a K-Means clustering approach within the MAPS software, in order to determine the total amounts of elements contained within these areas and the fraction of the cell that they cover. “Intracellular areas outside of storage regions” were defined as the areas within the boundaries of the cell, that did not have a high Cu or Fe content, respectively. They were identified as the inverse intracellular areas of the “areas containing storage compartments”.

Enrichment of elements within storage compartments (Figure 5A) was determined with two criteria. Elements within “storage compartments” were required to accumulate to significantly greater levels than they would naturally, given the total cellular content of the element and the fraction of the area of the cell that contain storage compartments. 9.9 (±3.8) % and 6.6 (±1.3) % of the area of the cell, on average, contained high concentrations of Cu or Fe, respectively, and therefore were considered to contain storage compartments. This fraction of the cell (average) is also presented by a black, dotted line in Figure 5A. The average and standard deviation of the fraction of the area containing storage compartments between individual cells, as well as the average and standard deviation of the fraction of the elemental content in areas containing storage compartments between individual cells was used for the statistical analysis. Since there might be a bias for areas in the center of the cell, where the cells thickness should be maximal, we also required individual elements to exceed the abundance of S in these areas, which was the element within our analysis that best correlated with biomass, was not affected by treatment or strain background and is most equally distributed throughout the cells (see Supplemental Figures 1-3), and therefore, within the parameters of the experiment, should best approximate the volume. The average S content in the areas containing storage compartments was determined between individual cells and was added to Figure 5A as a dashed line. In the areas that have high Fe/Cu content, we found between 12-23 % of the total cellular S. We recognize that this approach could be overly conservative with respect to identification of other elements, including S, that may be enriched within the storage organelles. But given the 2-dimensional limitations of the methodology, we aimed to raise the bar for elements to be considered sequestered. The “Intracellular areas outside of storage regions” were used to determine the intracellular concentration of Fe and Cu outside of the storage organelles (Figure 5C).

In addition to the content of the different areas within the cell, the number of storage compartments / cell was determined manually by counting individual secluded areas within the cells. This number, together with the total area containing storage compartments, was used to determine the average dimensions (area and radius) of the storage compartments within individual cells (Figure 5D).

## Author contributions

S.S. and S.S.M. conceptualized the project. S.S., D.S. and C.H. prepared samples for elemental analysis, S.C., M.R., S.S., D.S. and C.H. generated elemental distribution maps at the synchrotron. S.S., M.R. and S.C. performed advanced XFM image analysis. S.S. generated the ICP MS-MS and TOC/TN bulk cell data. S.S. and S.S.M. wrote the manuscript draft, all authors reviewed and edited the paper.

## Data and materials availability

All data are available in the main text or the supplementary materials.

