## Supplemental Figures for "Single-cell Visualization and Quantification of Trace Metals in Chlamydomonas Lysosome-Related Organelles"

### Supplemental Figure 1

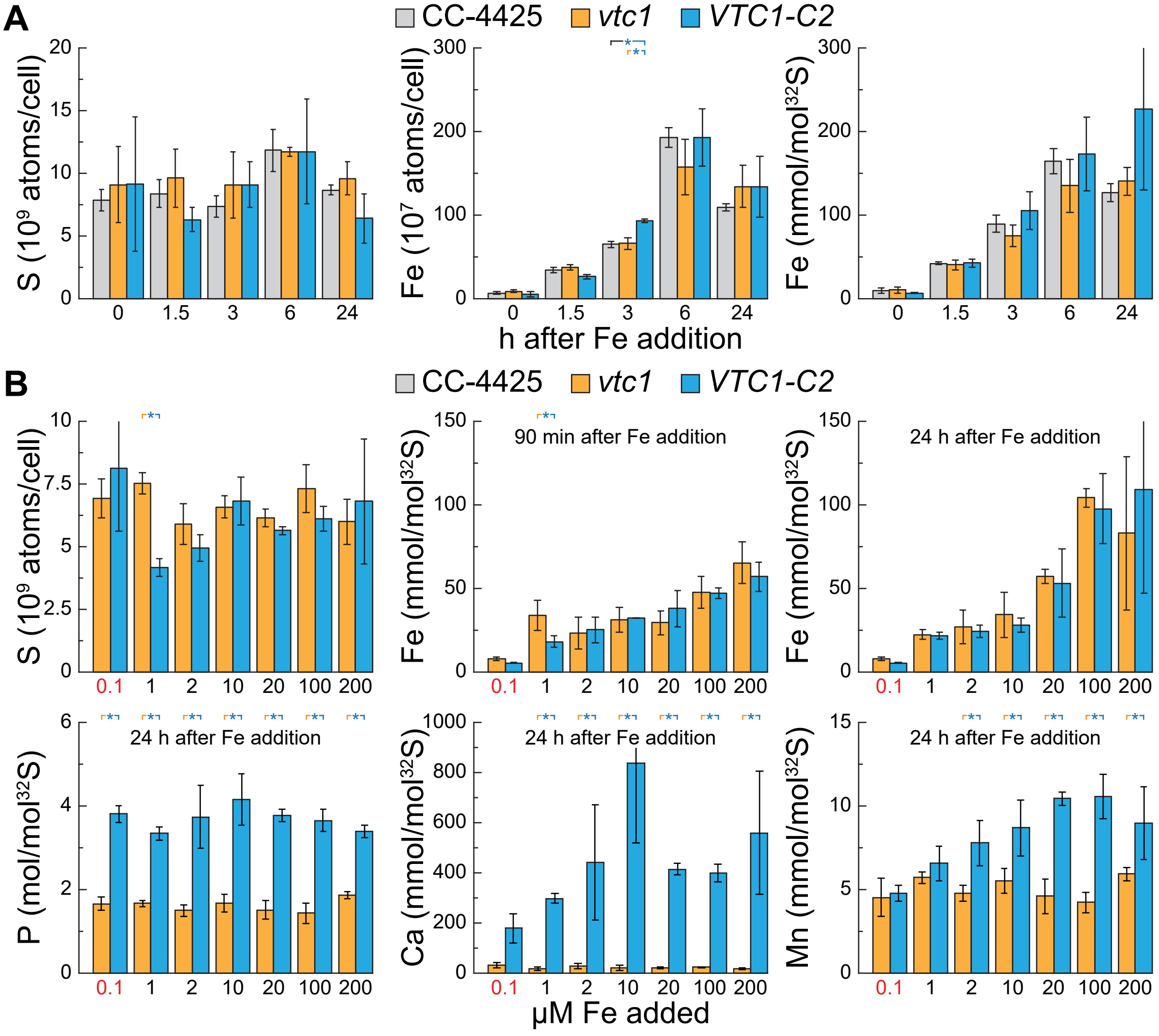

**Supplemental Figure S1: S is a good reference element in Fe variation experiments. Luxury Fe uptake is not affected in *vtc1* mutants**

(A) Comparison between the total S and Fe content per cell (left, middle), and Fe normalized to the cellular S content (right) of *Chlamydomonas reinhardtii* wild-type CC-4425 (grey bars), the *vtc1* mutant (orange bars) and complemented strain (VTC1-C2, blue bars) as measured by ICP-MS/MS, either in Fe-limiting conditions (0h, 0.1 µM Fe) or as a function of time (1.5, 3, 6, and 24 h) after the addition of 200 µM Fe to Fe limited cells. Error bars indicate standard deviation of n≥3 individual cultures. Asterisks indicate significant differences (ANOVA with subsequent means comparison (Fisher LSD), $P \leq 0.05$).

(B) Total cellular S, Fe, P, Ca and Mn content of the *vtc1* mutant (orange bars) and complemented strain (VTC1-C2, blue bars), as measured by ICP MS/MS either in Fe-limiting conditions (0.1 µM Fe, before Fe addition) and as a function of back-added Fe (1, 2, 10, 20 (replete), 100 and 200 µM in the growth media) 90 min or 24 h after the addition to Fe limited cells. S content is normalized per cell (left, upper panel) and compared to Fe, P, Ca and Mn normalized to total cellular S content as a measure for total biomass (middle, right, upper panel and all of the lower panels). Error bars indicate standard deviation of n≥3 individual cultures. Asterisks indicate significant differences (ANOVA with subsequent means comparison (Fisher LSD), $P \leq 0.05$).

**Supplemental Figure 2**

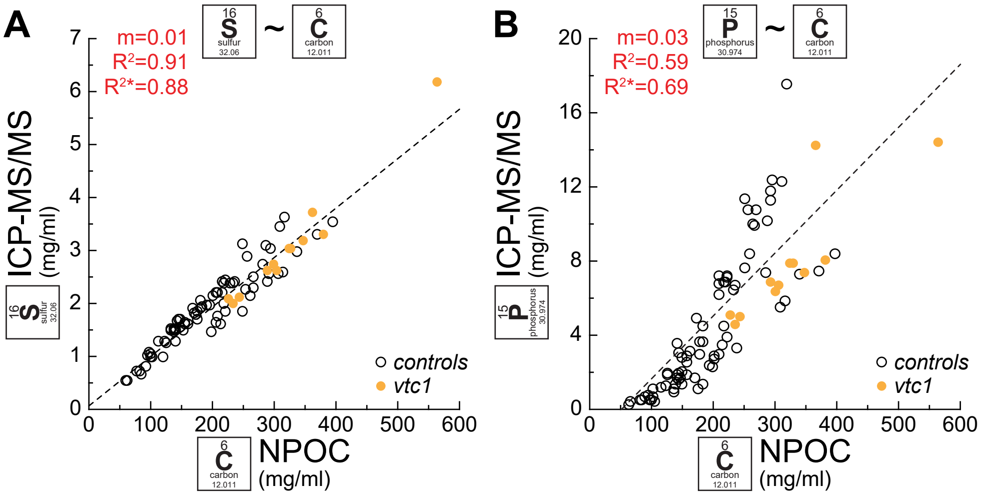


**Supplemental Figure S2: S correlates well with biomass**

(A/B) Correlation of biomass, measured as Non-Purgeable Organic Carbon (NPOC) on a Total Organic Carbon and Total Nitrogen analyzer (TOC/TN, x-axis) and total cellular S (A) and P (B) content as measured by ICP-MS/MS (Oxygen mode, y-axis). Each individual point corresponds to an individual sample from experiments upon Fe addback to Fe limited cells (Fe fast transition, Figure 2 and Figure S1). Black outlined circles indicate control samples with wild-type levels of VTC1 (CC-4425 and *VTC1-C2*), orange filled circles indicate samples from the *vtc1* mutant, which has reduced cellular P content. $R^2$ and slope (m) of a linear regression analysis (black dashed line) using all samples are given in the top left corner, $R^{2*}$ was calculated similarly using only control samples. All linear regression calculations were performed in OriginPro (v9.1).

**Supplemental Figure 3**

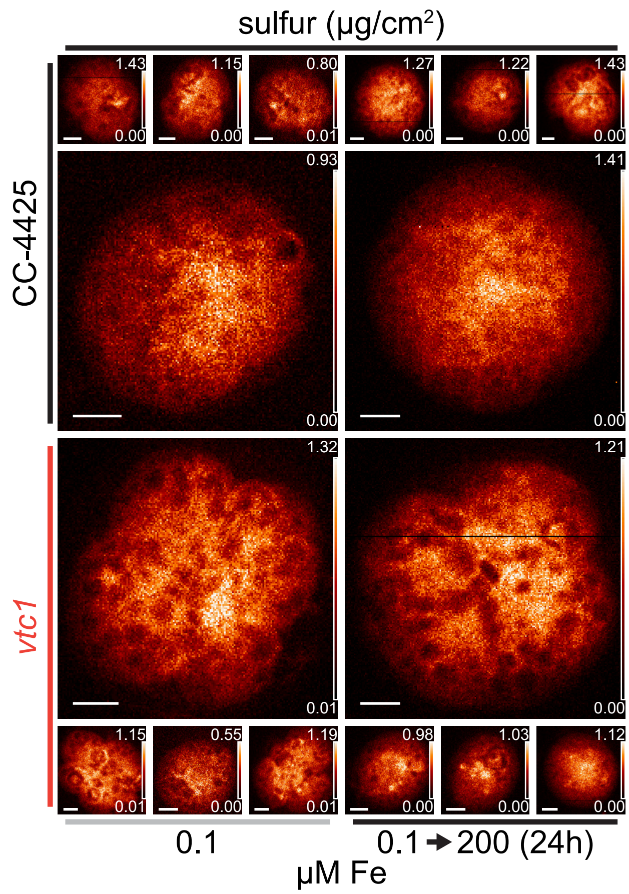


**Supplemental Figure S3: S distribution in Fe accumulating cells**

S distribution in 4 individual cells of the Chlamydomonas wild-type (CC-4425) and the *vtc1* mutant, in Fe-limited or 24 h of Fe–excess conditions (from Figure 2B), as measured by X-ray fluorescence microscopy at the Bionanoprobe (APS). Images of chemically-fixed alga cells show S distribution ranging from minimum (black) to maximum (orange) concentration in µg/cm$^2$. Each cell was scaled individually, actual minimum and maximum concentrations for each cell are denoted in the lower and upper right hand corner of each image, respectively, white scale bars in the lower left corner of the images indicate 2 µm size. Cells were prepared as described in Fig1B.

**Supplemental Figure 4**

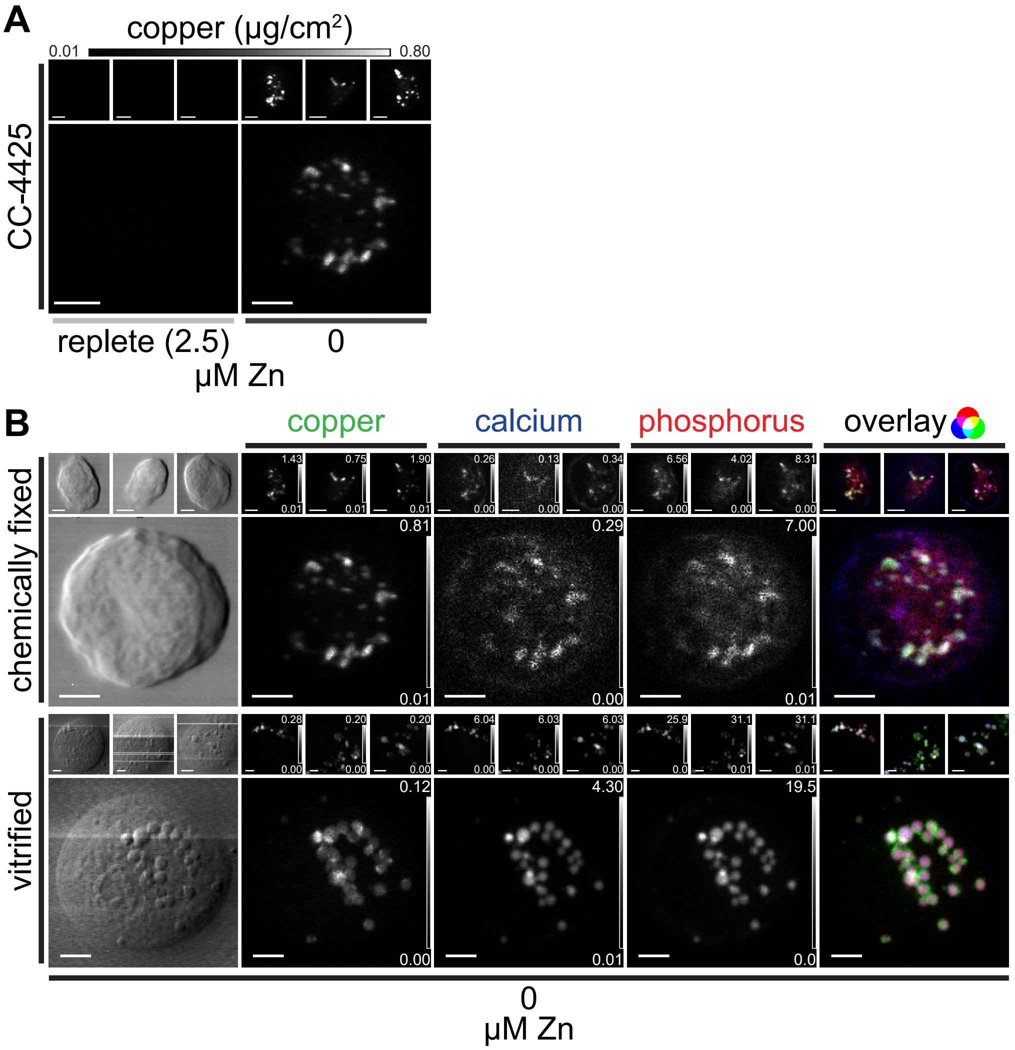

**Supplemental Figure S4: Cu distribution and co-localization in Zn deficiency**

A: Cu distribution in 4 individual CC-4425 wild-type cells, in Zn-replete (left) or Zn-deficient (right) conditions (from Figure 1B), measured by X-ray fluorescence microscopy at the Bionanoprobe (APS). Cu distributions for all cells are depicted between shared, fixed minimal (black) and maximal (white) Cu concentrations, denoted above the two leftmost image sets (in µg/cm$^2$), scale bars in the lower left corner indicate 2 µm.

B: For the same Zn-deficient cells, individually scaled minimum and maximum concentration of Cu, Ca and P for each cell are denoted in the lower and upper right hand corner of each image, scale bars in the lower left corner indicate 2 µm. The cells were chemically fixed and stored at room temperature (upper panel) and are compared to 4 individual cells in the same condition (Zn-deficiency) that were rapidly vitrified (lower panel), the images were acquired using fly-scan mode (continuous motion in the horizontal direction) at low temperatures with 70 nm step size and 300 ms dwell time per position. The fluorescence maps were created by performing peak area fitting for every position utilizing the MAPS software suite. The rightmost image shows the overlay of the three elements, where Cu contributes the green, Ca the blue and P the red color.

**Supplemental Figure 5**

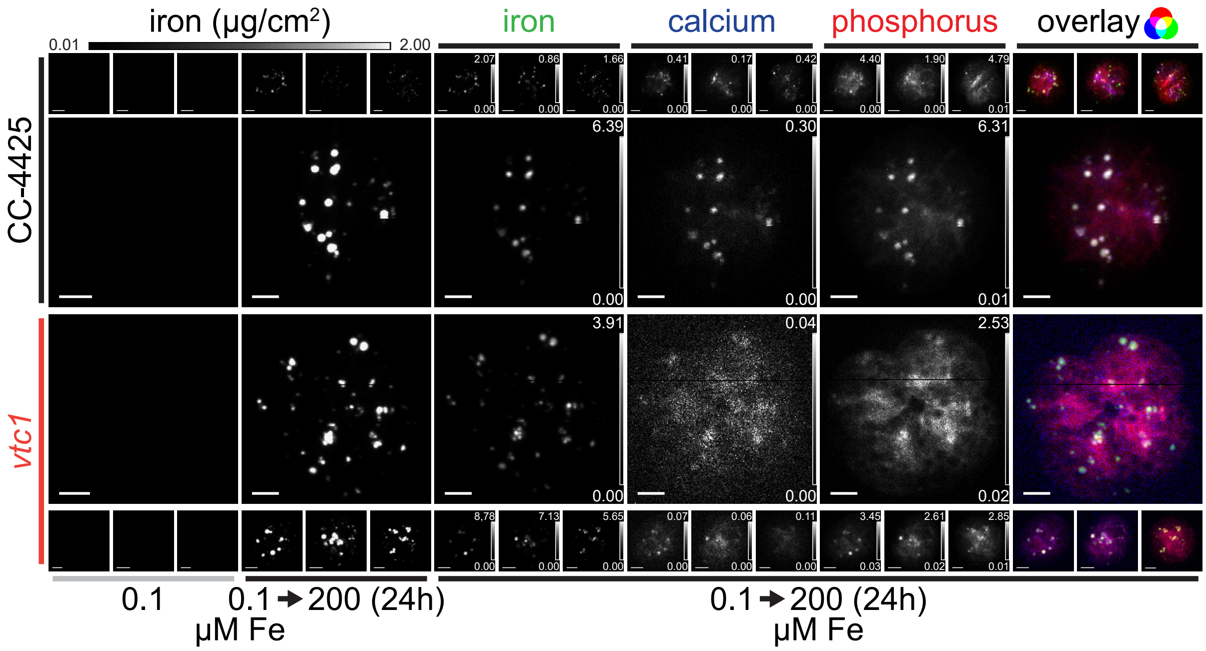


**Supplemental Figure S5: Fe distribution and co-localization in Zn deficiency**

Fe distribution in 4 individual CC-4425 wild-type and *vtc1* mutant cells, in Fe-limited conditions or after 24 h of Fe–excess (from Figure 2B/C), measured by X-ray fluorescence microscopy at the Bionanoprobe (APS). Fe distributions for all cells are depicted between shared, fixed minimal (black) and maximal (white) Fe concentrations, denoted above the two left-most image sets (in µg/cm$^2$). For the same cells, individually scaled minimum and maximum concentration of Fe, Ca and P for each cell are denoted in the lower and upper right hand corner of each image in the middle and on the right of the panel, respectively, scale bars in the lower left corner indicate 2 µm. The cells were fixed and stored at room temperature, the images were acquired using fly-scan mode (continuous motion in the horizontal direction) at low temperatures with 70 nm step size and 300 ms dwell time per position. The fluorescence maps were created by performing peak area fitting for every position utilizing the MAPS software suite. The rightmost image shows the overlay of the three elements, where Fe contributes the green, Ca the blue and P the red color.

**Supplemental Figure 6**

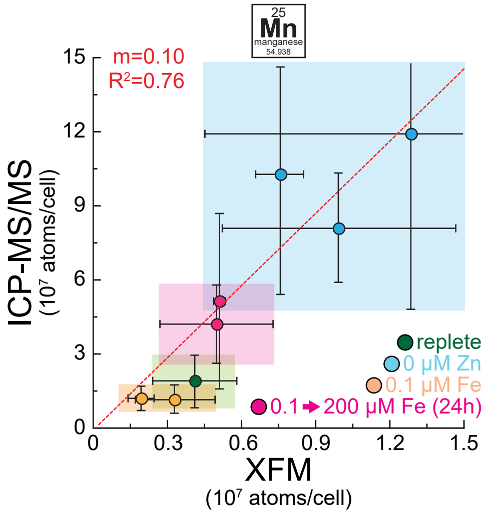


### Supplemental Figure S6: Correlation of Mn content between XFM and ICP-MS/MS

Correlation of cellular Mn content as measured by X-ray fluorescence microscopy (x-axis) and ICP-MS/MS (y-axis). Error bars in x and y direction indicate standard deviation in the measurements between individual cells (XFM) or between at least between 3 experiments (ICP-MS/MS). Cyan colored areas indicate Zn-deficient conditions (5-fold Mn accumulation), magenta areas indicate 24h of Fe-excess (two-fold Mn accumulation), orange areas indicate Fe-limited conditions (low Fe, medium Cu) and green areas indicate replete conditions. $R^2$ and slope (m) of a linear regression analysis (red dotted line) are given in the top left corner and were calculated using OriginPro (v9.1). Each individual dot represents a different experiment or strain, measured within the respective conditions at the APS.
